# Meso2EM: a cross-scale CLEM workflow linking mesoscale functional imaging to targeted electron microscopy

**DOI:** 10.64898/2026.08.25.746890

**Authors:** Ikumi Oomoto, Motohide Murate, Jaerin Sohn, Masaru Tamura, Sayuri Hatada, Naomi Egawa, Maya Odagawa, Mitsuo Suga, Yasuo Kawaguchi, Masanori Murayama, Yoshiyuki Kubota

## Abstract

Meso2EM is a correlative light and electron microscopy workflow that transfers neurons selected from mesoscale functional images to targeted electron microscopy. We recorded Ca²⁺ signals from layer 2/3 neurons across a contiguous 3 × 3 mm cortical field in awake mice and reidentified a selected neuron after fixation and tangential sectioning. Lectin-labeled vascular architecture served as a shared landmark across in vivo two-photon imaging, confocal microscopy, laboratory micro-CT of resin-embedded tissue, and block-surface scanning electron microscopy, guiding focused-ion-beam scanning electron microscopy to the target cell body. The same progressive-targeting principle also supported serial ATUM-SEM reconstruction of an in vivo–tracked dendrite and serial transmission electron microscopy of optically selected dendrites from a patch-clamp-recorded Martinotti cell. Meso2EM therefore provides a practical route for preserving target identity across large changes in scale and specimen state while restricting electron-microscopy acquisition to a selected region.

## Introduction

Advances in *in vivo* wide-field two-photon microscopy now enable the recording of neuronal activity across multiple cortical areas spanning millimeters at single-cell resolution^1–4^. Such mesoscale functional imaging not only reveals coordinated activity across areas that is difficult to observe from recordings restricted to local circuits, but also enables analysis of individual neuronal activity in relation to spatial position within a network. Indeed, analyses of functional networks constructed from single-cell activity reveal spatially intermixed modules and neuronal populations with different node degrees^5,6^. Determining the dendrites, synapses, intracellular structures, and local circuit environments of specific cells identified within mesoscale images is therefore important for linking large-scale activity to cellular structure.

Electron microscopy (EM) visualizes synaptic connections, membrane structures, and intracellular organelles that cannot be resolved by light microscopy. Studies combining in vivo functional recording with serial EM have demonstrated that activity properties and local connectivity can be related in the same cells^7,8^. However, imaging an entire tissue area of several square millimeters at high EM resolution merely to locate a target is impractical in terms of both acquisition time and data volume^9^. What is needed is a method that preserves the identity of cells selected from wide-field data, progressively narrows the search space, and accurately positions a limited EM volume over the target cell.

Correlative light and electron microscopy (CLEM) has developed to link functional and optical information with ultrastructure in the same specimen^10–12^. In neural tissue, diverse approaches have correlated blood vessels, nuclei, fluorescent labels, tissue contours, and artificial marks to guide target cells or neuronal processes to serial EM^13–17^. The use of X-ray microscopy or micro-computed tomography (micro-CT) as a nondestructive intermediate image of resin-embedded blocks to define sites for volume EM is also well established^13–16^. Moreover, a previous study integrating two-photon calcium imaging, synchrotron X-ray microtomography, and volume EM demonstrated that functional information and three-dimensional ultrastructure can be linked across scales^13^. Nevertheless, a practical route for guiding a specific neuron from cell-resolution functional images spanning several millimeters, through laboratory-based micro-CT, to targeted FIB-SEM has not been fully established as an integrated workflow.

Here, we developed and demonstrated Meso2EM (Meso-to-EM), a workflow linking mesoscale functional imaging to targeted EM. Its principal design features are to preserve orientation between the in vivo imaging plane and the post-fixation sectioning plane, acquire landmarks visible across multiple modalities such as vascular architecture, and iteratively narrow the search space through confocal microscopy, micro-CT, and block-surface SEM. In the core application, a single neuron recorded by two-photon calcium imaging over a 3 × 3 mm field of view was identified in the FIB-SEM milling face after tangential sectioning, confocal microscopy, and micro-CT. To demonstrate the broader applicability of this design principle, we also present ATUM-SEM analysis of a dendrite tracked in vivo and serial TEM analysis of a patch-clamp-recorded cell. The aim of this study was to define and demonstrate a continuous experimental pipeline for transferring cells selected in wide-field functional images to targeted EM and to define its key implementation steps.

## Results

### Overview of the Meso2EM workflow

In Meso2EM, a target identified in the initial optical image was not converted directly into EM coordinates; instead, it was progressively reidentified using landmarks shared between successive stages (Fig. 1, Supplementary video 1). During transfer from in vivo wide-field two-photon images to fixed tissue, we used the shape of the cranial window, tissue contours, needle-puncture marks, and large blood vessels. During transfer from fixed sections to the resin-embedded tissue block, we used the three-dimensional trajectories of lectin-labeled blood vessels. After resin embedding, resin-filled vascular lumens, which exhibited relatively low X-ray absorption compared with heavy-metal-stained neural tissue, were visualized by micro-CT and correlated with vascular cross-sections observed by block-surface SEM. This process narrowed the millimeter-scale search area to a local region accessible by FIB-SEM.

**Figure 1.**
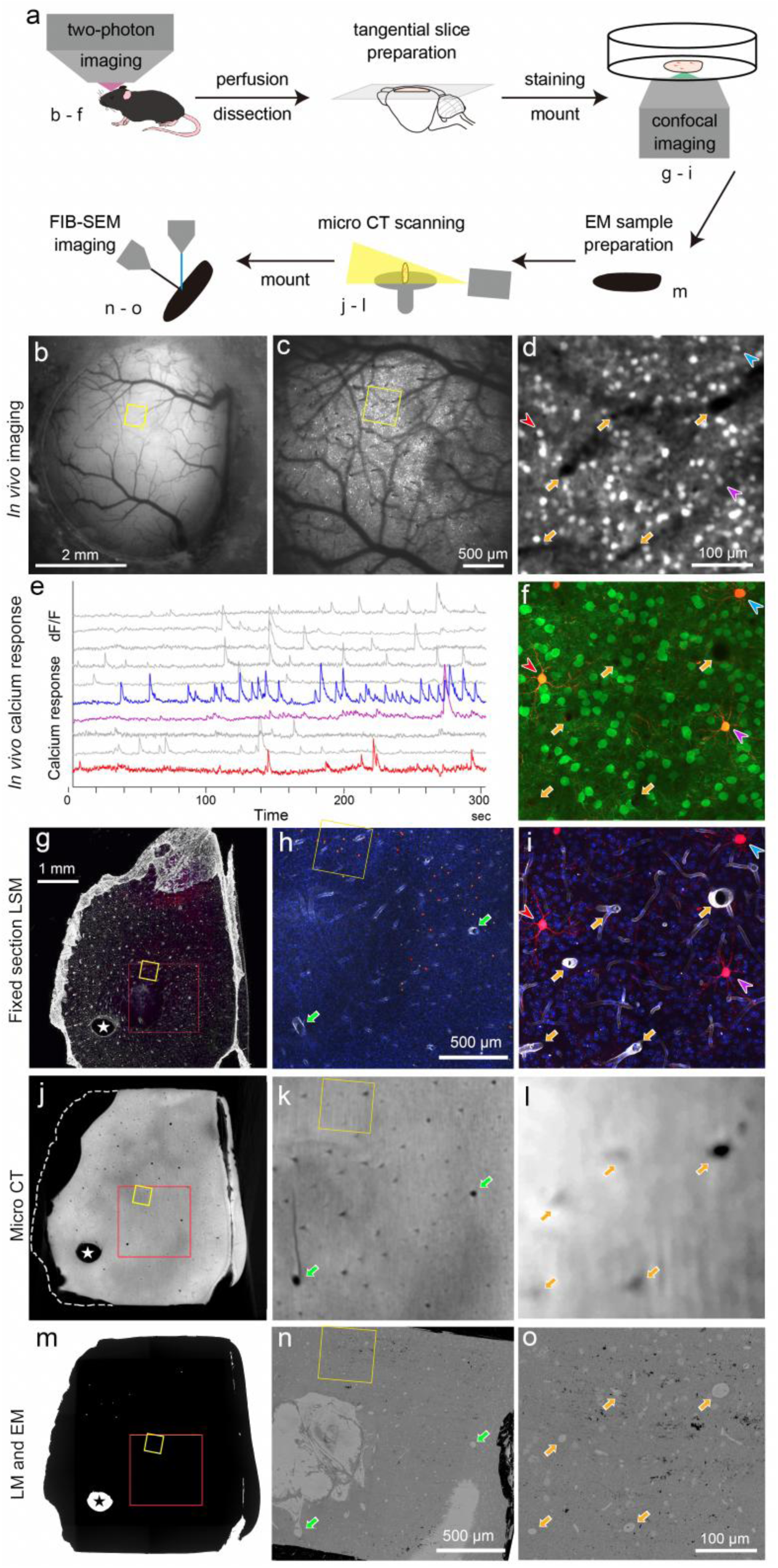
**(a)** Schematic pipeline of CLEM with in vivo calcium imaging in a large area and focus ion beam electron microscopy. Panel labels (a–o) in each cartoon indicate the corresponding panels. **(b)** This is a dorsal view of a mouse that has been implanted with a cranial window for wide-field two-photon imaging. The yellow square indicates the ROI for a subsequent EM study. **(c)** A snapshot from two-photon calcium imaging of cortical layer 2 neurons labeled with GCaMP7.09 in a 3 mm x 3 mm field of view (FOV). The yellow square indicates the ROI for the subsequent EM study. This area is the cropped region for d. **(d)** An enlarged image of part of the FOV. The red arrowhead indicates the neuron of interest for CLEM in this study. Yellow arrows indicate blood vessel holes. The blue and violet arrowheads indicate neurons for activity tracing in e. **(e)** Representative spontaneous Ca²⁺ signals from ten randomly selected neurons in the area shown in d. The red trace indicates the Ca²⁺ signal from the neuron of interest in this study. **(f)** A confocal image of a wet tangential brain slice horizontal to the 2P imaging plane. The same neurons were observed in the brain slice. Green: G-CaMP7.09; red: tdTomato. **(g)** A confocal image of a wet tangential brain slice stained with Lectin-647 to visualize blood vessels. A red square indicates the area for h, and a yellow square indicates the ROI for the later EM study in H. **(h)** A maximum intensity projection image of z-stacked confocal images of the tangential brain slice. Green arrows indicate the large blood vessels used to align the three different modality images. **(i)** A maximum intensity projection image of z-stacked confocal images of the tangential brain slice. The DAPI signal (blue), the tdTomato signal (red), and the Lectin647 signal (white) are shown. **(j-l)** An image from serial micro-CT scanning of the tangential brain slice after EM sample preparation. The left side of the tissue block shown with the dotted line in I was cut out to allow for better imaging with the micro-CT. **(m)** A bright-field image of the tangential brain slice after EM sample preparation using the rOTO protocol. This is the same tissue block shown in F and I. **(n)** A surface image of the sample obtained by SEM before FIB-digging. **(o)** An enlarged view of the yellow square in m and n. **(n)** Images in d, f, i, l, o, and g, j, m, and h, k, n shows the identical region captured using different microscopy in the same magnification.

The three implementations presented here differed in their optical inputs, intermediate landmarks, and EM endpoints, but shared the principle of verifying target identity across multiple stages (Extended Data Table 1). We first describe the core Meso2EM application linking wide-field functional imaging, micro-CT, and FIB-SEM, followed by applications to dendrites and patch-clamp-recorded cells.

**Extended Data Table 1.** Information transfer and endpoints in three CLEM implementations.

| Implementation | Functional/optical input | Intermediate landmarks | EM endpoint | Outcome demonstrated in this study |
| --- | --- | --- | --- | --- |
| Wide-field functional imaging–FIB-SEM (core Meso2EM application) | Awake $3 \times 3$ mm two-photon $\text{Ca}^{2+}$ images; G-CaMP7.09; tdTomato; target-cell morphology | Cranial window; tissue contour; needle-puncture mark; large blood vessels; lectin-labeled vasculature; vascular lumens in micro-CT; vascular cross-sections at the block surface | Target cell body in the FIB-SEM milling face | The same cell recorded by wide-field functional imaging was localized within the resin block and reached during FIB milling |
| In vivo dendritic imaging–ATUM-SEM | Repeated two-photon images in Thy1-eGFP-M mice; dendritic and spine morphology | Dendritic branching; tissue contour; lectin-labeled vasculature; nuclei; vascular lumens in serial sections | Target dendrite, spines, and synapses | The dendrite tracked in vivo was reidentified in serial SEM images and reconstructed in three dimensions |
| Patch clamp–serial TEM | Whole-cell recording in acute slices; biocytin/DAB labeling; cell morphology | Dendritic branching; capillaries; DAB-positive processes; shapes of cells and tissue spaces | Selected dendritic segments and associated synapses | Optically selected Martinotti-cell dendrites were identified by serial TEM and reconstructed in three dimensions |

### Selection of a target cell from wide-field two-photon calcium imaging

G-CaMP7.09 was expressed in cortical layer 2/3 beneath a cranial window in awake mice, and a contiguous 3 × 3-mm field of view was imaged using two-photon microscopy (Fig. 1b–d)^4,5^. The field of view encompassed the somatosensory and motor cortices, as well as parts of higher-order cortical areas, allowing spontaneous Ca²⁺ signals to be recorded from numerous neurons. A Ca²⁺ trace was extracted from the neuron selected for subsequent CLEM analysis, confirming that it had been functionally recorded within the wide-field network (Fig. 1e). A subset of neurons, including the selected neuron, also expressed tdTomato, which enabled morphological reidentification after fixation (Fig. 1f).

The objective at this stage was not to test a causal relationship between a specific activity pattern and ultrastructure, but to determine whether the identity of a single neuron selected from a wide-field functional dataset could be preserved throughout the workflow and ultimately confirmed by EM. The target neuron was selected before fixation according to prespecified criteria encompassing its functional properties, fluorescence labeling or morphology, and the anatomical and technical feasibility of subsequent EM targeting. Neurons that did not satisfy these requirements were excluded. The neuron analyzed here met all of the prespecified criteria and was therefore selected for CLEM analysis.

### Reidentification of the same cell in post-fixation tangential sections

After two-photon imaging, the brain was perfusion-fixed and tangential sections were cut approximately parallel to the imaging plane. The shape of the cortical surface adjacent to the cranial window and needle-puncture marks placed outside the field of view preserved section orientation and position. Sections were stained with DAPI and fluorescently labeled lectin, and nuclei and vascular walls were acquired by confocal microscopy. Comparison of tissue contours, large blood vessels, vascular branch points, and the distribution of tdTomato-positive cells between the two-photon and confocal images enabled the target cell from the two-photon image to be reidentified in the fixed section (Fig. 1f–i). The confocal z-stack preserved the depth relationship between the target cell and surrounding vasculature and provided a three-dimensional map for the subsequent search within the resin block.

### Tracking vascular trajectories within the resin block by micro-CT

The section containing the target was heavy-metal stained and embedded in epoxy resin. Because direct observation of fluorescence became difficult after embedding, laboratory-based micro-CT was acquired as a nondestructive intermediate dataset (isotropic voxel size, 6.011 µm). Heavy-metal-containing neural tissue showed high X-ray absorption, whereas resin-filled vascular lumens appeared as relatively low-signal structures, allowing vascular trajectories within the block to be tracked in three dimensions (Fig. 2, Supplementary video 2). Large vessels and branching patterns at the section surface and within the tissue were compared between the confocal z-stack and the XY, XZ, and YZ planes of the micro-CT data, enabling the vascular arrangement adjacent to the target cell to be correlated within the resin block.

**Figure 2.**
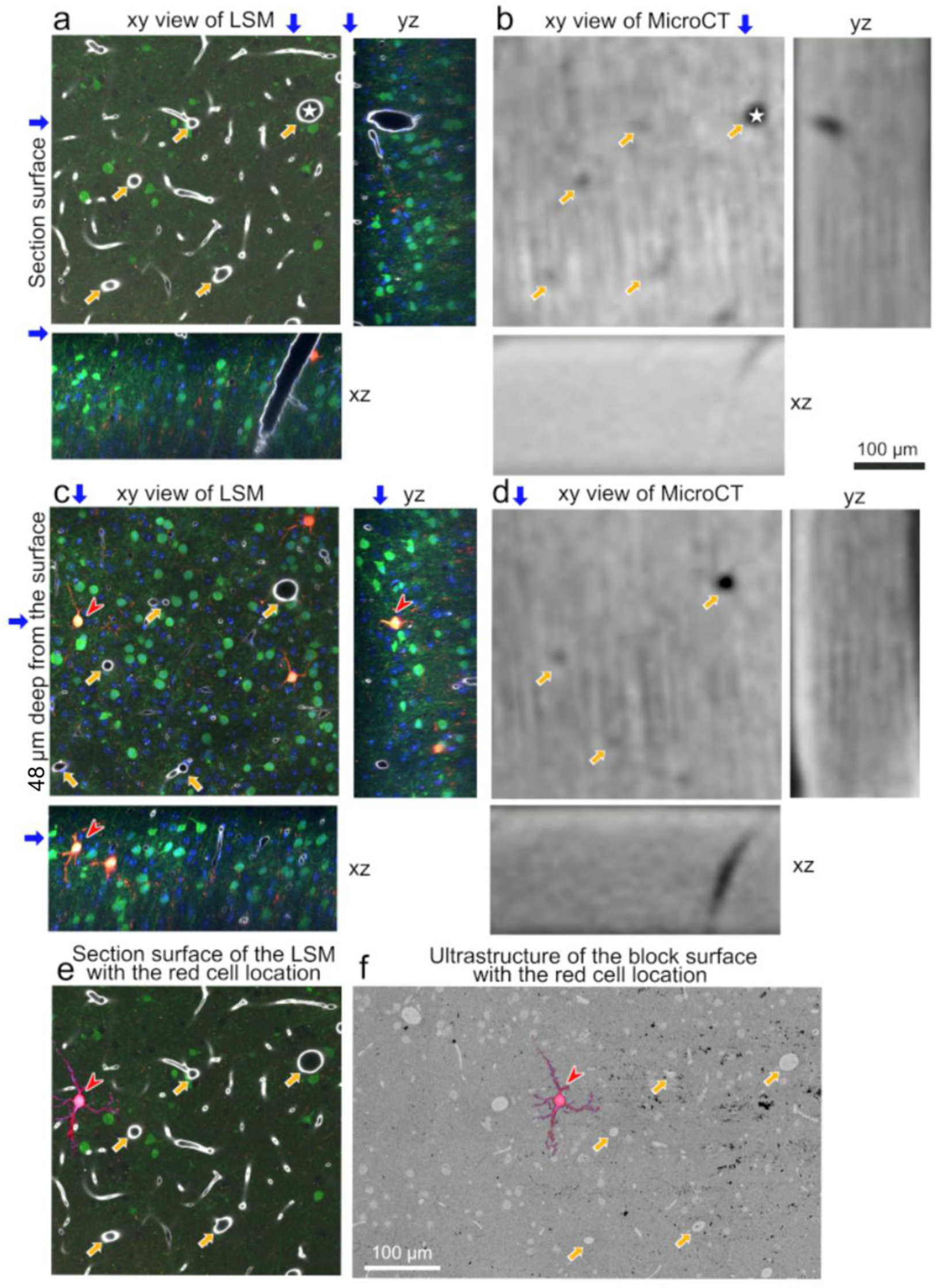
Image alignment of LMs and micro-CT data. **(a, b)** Micrographs of the section surface. The yellow arrows indicate the blood vessels that were used to align the two images. **(a)** XY, YZ, and XZ views of the confocal Z-stack images. Blue arrows indicate the positions of the orthogonal planes: arrows in the XY view indicate the positions of the YZ and XZ planes, whereas arrows in the YZ and XZ views indicate the corresponding depth of the XY plane. **(b)** Micro-CT data. **(c, d)** Micrographs at 48 µm below the surface in the wet brain slice. **(c)** XY, YZ, and XZ views of the confocal Z-stack images. A red arrowhead indicates the cell targeted in the FIB-SEM imaging portion. **(d)** Images of the micro-CT data corresponding to c, according to the blood vessel pattern. **(e)** The tdTomato-labeled target cell selected for FIB–SEM imaging, overlaid on the LSM image of the section surface shown in a. **(f)** The position of the target cell overlaid on the block-face image, with the corresponding capillaries matched to those shown in e.

At this stage, micro-CT did not serve to identify the cell body directly; rather, it acted as a coordinate bridge that preserved the cell–vessel relationship obtained by optical imaging after resin embedding. The block was trimmed with reference to vessel positions predicted from the micro-CT volume (Fig. 1j, m), and the corresponding vascular cross-sections were confirmed in block-surface SEM images (Fig. 1k, l, n, o). This defined the region to be milled by FIB-SEM and the depth range in which the target was expected to appear.

### Reaching the target cell body by FIB-SEM milling

After confirming vascular cross-sections in block-surface SEM images, FIB milling proceeded in the direction estimated from the confocal z-stack and microCT data (Fig. 2). During the search, the milling face was inspected at successive depths, and the arrangements of blood vessels, nuclei, and cell bodies were compared with the confocal images. The target cell body location in xy plane was estimated in relation to the landmarks such as capillaries (Fig. 2e, f). The depth of the cell was estimated with the LSM image data (Fig. 2c, d) and it appeared approximately 25 µm from the initial milling face and was tracked to approximately 38 µm (Fig. 3). Thus, a neuron recorded within a 3 × 3 mm in vivo functional image could be guided to the FIB-SEM milling face through vascular landmarks despite substantial specimen transformations during fixation, staining, and resin embedding.

**Figure 3.**
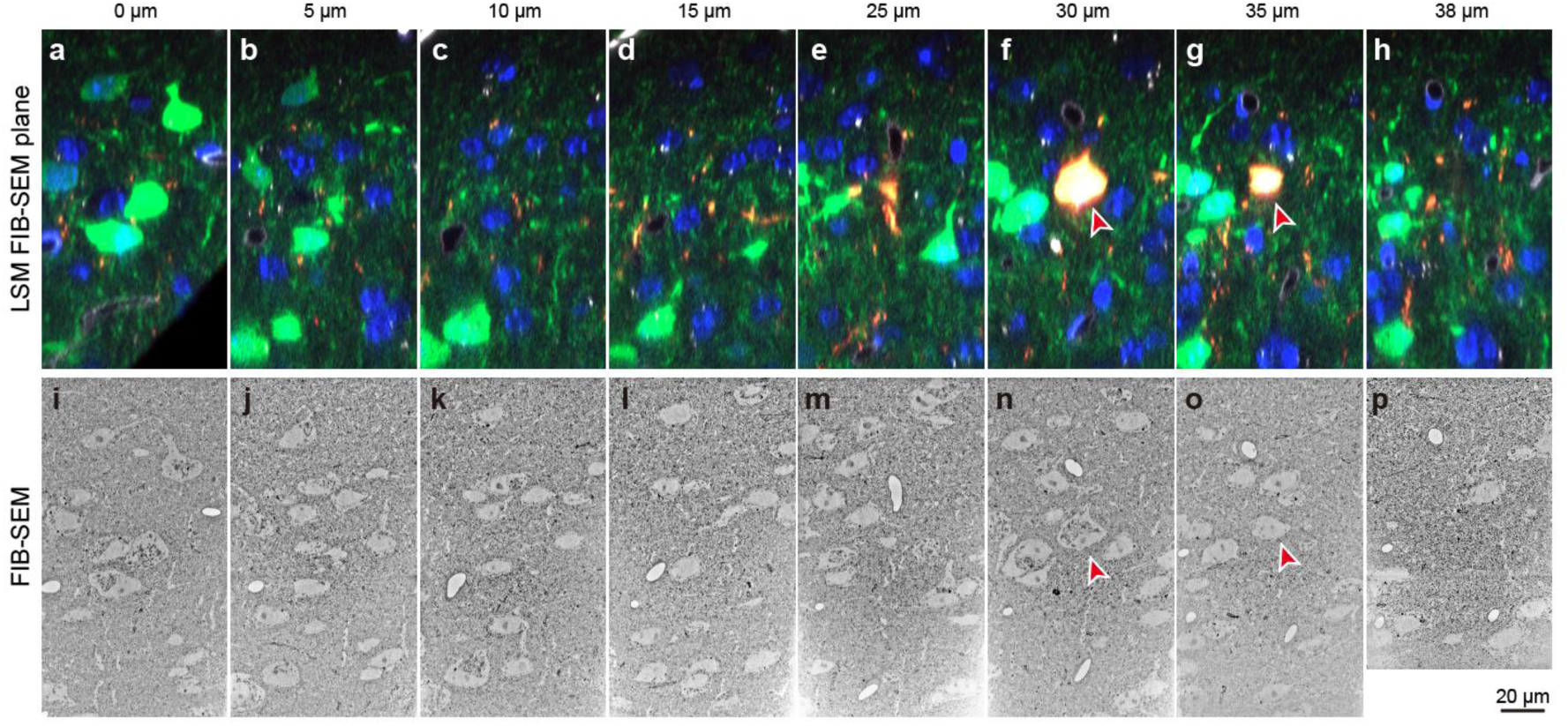
Correlated laser confocal micrographs and serial EM. **(a-h),** Cross sectional laser confocal micrographs including the ROI neuron (arrowhead, orange), G-CaMP7.09 labeled neurons (green), and DAPI stained nucleus (blue). (**i-p),** Electron micrographs showing equivalent regions as above confocal micrographs.

The EM endpoint demonstrated in this application was identification of the target cell body, rather than complete reconstruction of its entire axon, dendritic arbor, or synaptic connectivity. Nevertheless, the core Meso2EM route—progressively transferring the position of a specific cell from a wide-field functional image to the restricted acquisition volume of FIB-SEM—was demonstrated.

### Performance and reproducibility of the core Meso2EM route

In the core Meso2EM proof-of-concept experiment, identification of one target neuron was attempted and the same neuron was recovered at the FIB-SEM endpoint (1/1, 100% for this single attempt; Extended Data Table 2). At the 2P-to-confocal, confocal-to-micro-CT and micro-CT-to-FIB-SEM transfer steps, no appreciable positional discrepancy was observed using the anatomical and vascular landmarks available in the corresponding images. These observations were qualitative; localization distances were not measured separately at each step. The pre-trimming prediction nevertheless placed the target within the intended FIB-SEM search region.

The complete route required approximately 2 months, including preliminary optimization experiments. Critical steps and targeting decisions were discussed and jointly reviewed by several investigators, which supported consistent execution of the protocol when the specified procedure was followed. Although experiment 1 comprised only one CLEM preparation, experiments 2 and 3 implemented the same progressive-targeting principle through their respective pipelines, and in neither experiment did we encounter a case in which the intended target was lost before EM identification when the prescribed steps were followed. These observations support procedural reproducibility across the three implementations. Formal independent-animal and independent-operator reproducibility tests of the core route were not performed.

**Extended Data Table 2.**
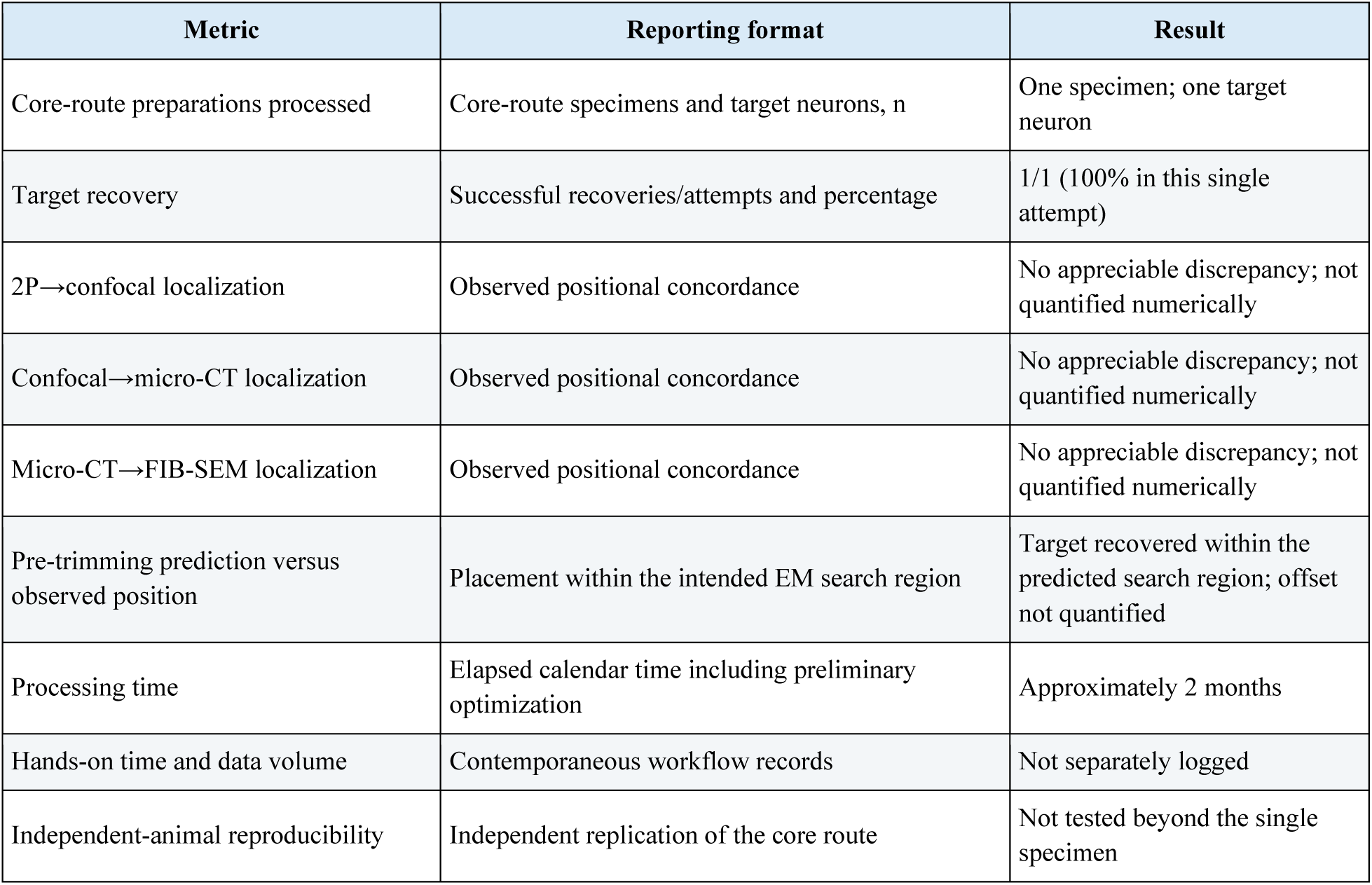
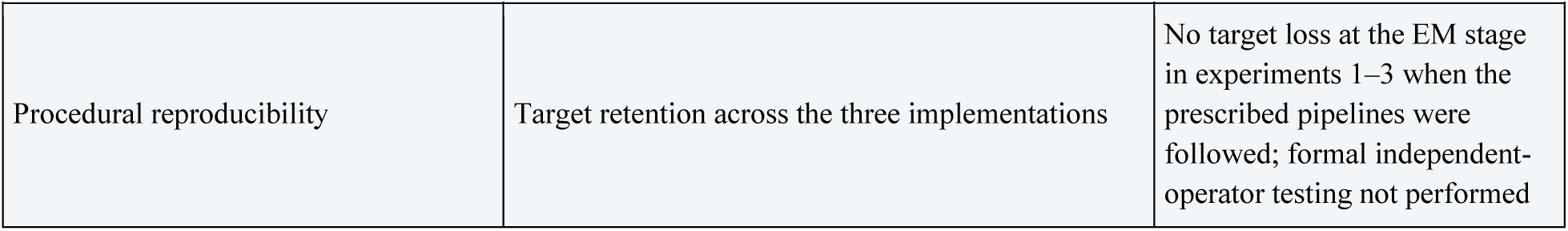
Observed performance and procedural reproducibility of the core Meso2EM route.

| Metric | Reporting format | Result |
| --- | --- | --- |
| Core-route preparations processed | Core-route specimens and target neurons, n | One specimen; one target neuron |
| Target recovery | Successful recoveries/attempts and percentage | 1/1 (100% in this single attempt) |
| 2P→confocal localization | Observed positional concordance | No appreciable discrepancy; not quantified numerically |
| Confocal→micro-CT localization | Observed positional concordance | No appreciable discrepancy; not quantified numerically |
| Micro-CT→FIB-SEM localization | Observed positional concordance | No appreciable discrepancy; not quantified numerically |
| Pre-trimming prediction versus observed position | Placement within the intended EM search region | Target recovered within the predicted search region; offset not quantified |
| Processing time | Elapsed calendar time including preliminary optimization | Approximately 2 months |
| Hands-on time and data volume | Contemporaneous workflow records | Not separately logged |
| Independent-animal reproducibility | Independent replication of the core route | Not tested beyond the single specimen |
| Procedural reproducibility | Target retention across the three implementations | No target loss at the EM stage in experiments 1–3 when the prescribed pipelines were followed; formal independent-operator testing not performed |

### Reidentification of a dendrite tracked *in vivo* by ATUM-SEM

We next applied the workflow to dendritic branches monitored longitudinally *in vivo.* Layer 5 pyramidal neurons in motor cortex were imaged through chronic cranial windows under a seed-grasp motor task training paradigm^17^. A cranial window was created above the primary motor cortex (M1) of a Thy-1-M line mouse, in which the cortical layer 5 pyramidal cells express green fluorescent protein (GFP) (Fig. 4a, b). We observed the arborization of tuft dendrites of layer 5 pyramidal cells under the 2-mm-diameter cranial window using two-photon laser microscopy (Fig. 4c–e) and analyzed the dynamics of spines on dendritic segments. After eight days of training and imaging, we perfused the mouse. The tissue sections must be cut in the same plane as the two-photon laser microscopy (Fig. 4f). The cortical region beneath the cranial window, where the round cranial window cover slip was clearly visible on the cortical surface (Fig. 4b, h, k), was roughly trimmed with a razor. The cerebral cortex surface beneath the cranial window was positioned upright on a glass plate. Then, an agar solution precooled to approximately physiological temperature was poured over the tissue and allowed to solidify at ambient temperature. We trimmed the agar block surface to be parallel to the underlying glass plate. Then, we inverted the block and mounted it onto a vibratome to section it into 300 µm-thick slices (Fig. 4f, g). The first tissue slices contained most of the dendritic segments observed under two-photon laser confocal microscopy, and the round cranial window was occasionally visible (Fig. 4h, k). The slice was incubated with Dylight 594 lectin to stain the capillary walls red and with DAPI to stain the nuclei blue for a few hours to mark landmarks for CLEM analysis (Fig. 4h-j). The slice was then observed under laser confocal microscopy. Using the overall shape of the slices and the large blood vessels as guides, we successfully identified the dendritic segments that had been imaged with in vivo 2-photon microscopy. Serial focus images were captured at 0.85 µm z-steps to include all dendritic segments. The total thickness of the dataset is about 200 µm. The light micrographs at different focus levels were meant to correspond to the EMs, and the anatomical landmarks were clearly identified in LMg and EM. Then, the slice was processed for electron microscopy (EM) and embedded in resin (Fig. 4k). Overall, the shapes of the wet tissue section and the tissue block section for EM were similar, but uneven shrinkage was observed. We estimated the portion where the dendritic segments were located (region of interest, ROI; Fig. 4h, k). The ROI region was cut out and trimmed for ultrathin sections. A total of 658 ultrathin sections, each 50 nm thick, were cut and collected on carbon nanotube-coated PET tape using an automated tape-collecting ultramicrotome (ATUM). The tape was then cut into strips and glued onto a 10 cm wafer using conductive double-coated adhesive tape. Ribbons of serial ultrathin sections were captured at low magnification (∼400 nm/pixel; Extended Fig. 3). The ultrastructure in the EMs was sufficient to identify the ROI by referring to the shapes and locations of the capillaries and blood vessels (Fig. 4i, j, l, m). After identifying the ROI, we imaged higher-resolution sections serially to locate the dendritic segments observed during in vivo imaging (Fig. 4m and n). We then reconstructed the segments for further analysis, including measuring spine volume, spine density, and synapse junction area (Fig. 4o)^17^.

**Figure 4.**
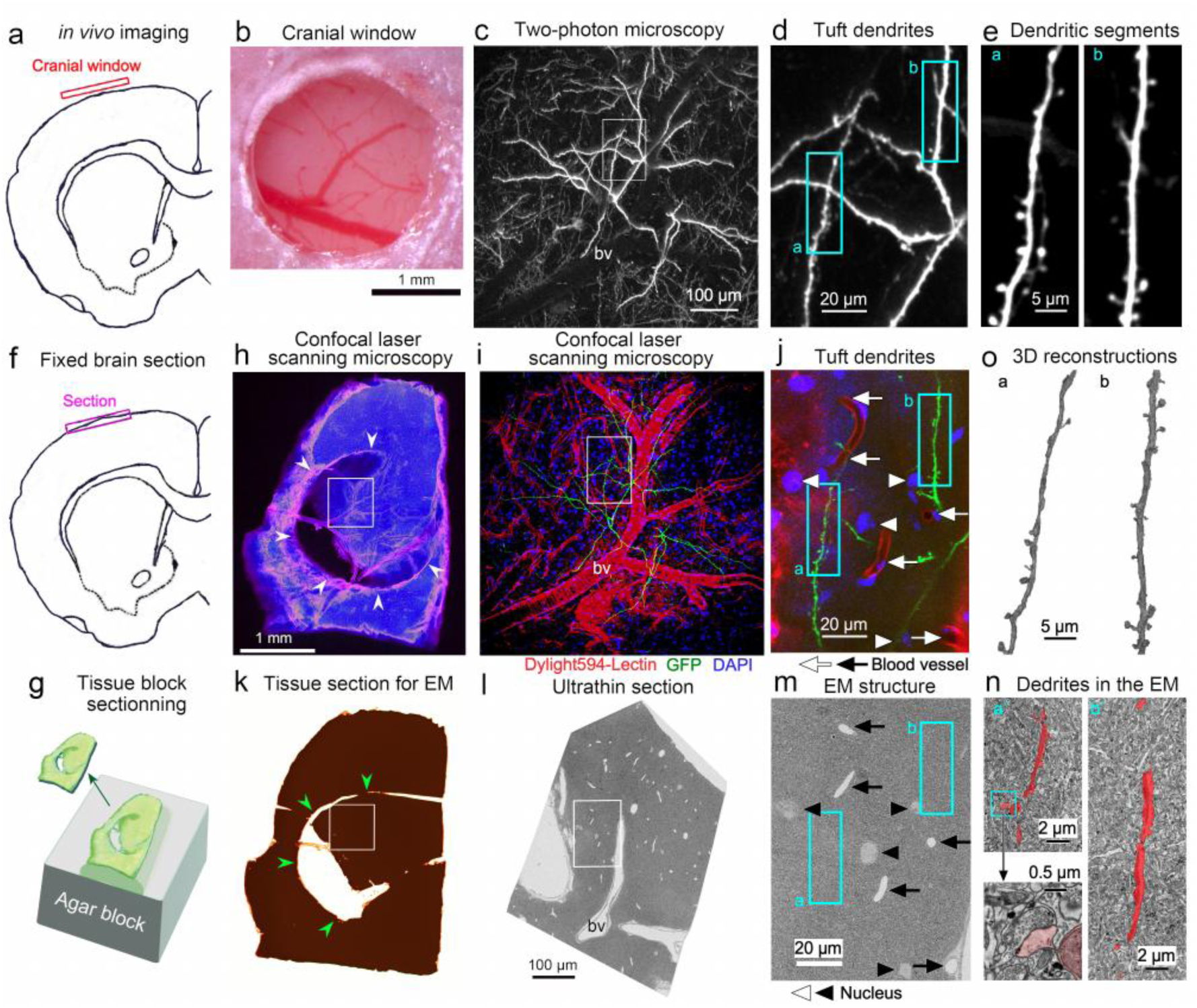
CLEM with in vivo imaging, confocal laser scanning microscopy and array tomography with scanning electron microscopy. **(a),** A schematic drawing shows the location of the cranial window on the primary motor cortex of the mouse in a coronal section. **(b)**. Cranial window above the primary motor cortex of the mouse. **(c)**. GFP-labeled tuft dendrites are observed using in vivo imaging with a two-photon microscope. bv: blood vessels. **(d**, **e)**: Enlarged views of the GFP-labeled dendritic segments shown in rectangles c and d, respectively. **(f)**. A schematic drawing showing a section with tangential cuts that efficiently include layer 1 of the mouse primary motor cortex at an identical angle to the cranial window. **(g)**. A schematic drawing shows how to obtain the first section from the surface of a fixed brain using an agar-embedded block. **(h)**. Laser confocal micrograph of the surface section directly beneath the cranial window. Arrows indicate marks left by the cranial window. Blue: DAPI; pink: blood vessels. **(i)**. An enlarged laser confocal micrograph of the square in h showing tuft dendrites (green), as seen in c. Blue: DAPI; pink: blood vessels. **(j)**. Enlarged laser confocal micrograph of the rectangle in i, showing two dendritic segments (green). Arrows indicate capillaries, and arrowheads indicate DAPI-stained nuclei. Blue: DAPI; pink: blood vessels. **(k)**. The surface of the brain tissue section for electron microscopy (EM) embedded in resin. This section is the same as the one shown in h. The arrows indicate the marks left by the cranial window. **(l)**. An electron micrograph showing the same blood vessel (bv) as in i. **(m)**. An enlarged electron micrograph of the rectangle in l, which is the correlated region shown in the laser confocal micrograph in j. Arrows indicate capillaries, and arrowheads indicate nuclei. **(n)**. Enlarged electron micrographs showing two dendritic segments (red) in rectangles a and b in m. The left bottom inlet shows synaptic input on the spine head. **(o)**. Reconstructed dendritic segments identical to those identified by in vivo imaging with a two-photon microscope. It is modified from Supplementary figure 12 in Sohn et al. (2022)

### Serial TEM reconstruction of selected dendrites from a patch-clamp-recorded cell

Finally, to examine the relationship between cellular physiology and ultrastructure, we analyzed a nonpyramidal cell that had been whole-cell recorded and injected with biocytin in an acute slice of rat frontal cortex^18–23^. Biocytin-filled neurons were visualized after DAB conversion and Neurolucida was used to reconstruct DAB-stained layer II/III Martinotti cells and the other non-pyramidal cells in the rat frontal cortex after patch clamp recording (see Fig. 5a and Extended Figure 4)^22^. The reconstruction is available at NeuroMorpho.Org (http://neuromorpho.org/neuroMorpho/index.jsp). A dendrogram was also obtained (see Extended Figure 4c). For three-dimensional reconstruction of the dendritic segment with serial electron micrographs (3DEM), we chose segments without bifurcations every 50 µm along dendrites that lacked artificially cut ends (see Extended Figure 4c). The corresponding segments were labeled in the serial focus light micrographs (LMgs, Extended Figure 4d). The Martinotti cell was stained with DAB, appearing blackish under light microscopy in a flat resin-embedded preparation. It could then be cut out from the tissue section under binocular microscopy and trimmed for ultrathin sectioning (see Extended Figures 4a and 4b). The target dendritic segments identified in the LMgs were located in the EMs using landmarks such as capillaries and electron-dense stained dendrites for reference (Fig. 5c and Extended Figures 4b-e). Dendritic segments stained black under EM observation could be observed in many serial ultrathin sections and were thus good landmarks for estimating the location of the target dendritic segment. Occasionally, the spine fragments of the target dendritic segment were also useful for this purpose. Overall, identifying the exact portion of the dendritic segment by looking at only the dendritic profile and nearby landmarks in a single 50-nm-thick ultrathin section under electron microscopy (Extended Figure 4e) is difficult. However, observing serial ultrathin sections allows us to estimate the portion of the dendritic segment more accurately than looking at only a single section image. Serial EMs containing stained neuronal profiles were manually imaged with extra care to cover the areas where black elements of dendritic segments were found in almost all serial ultrathin sections (Fig. 5d–g). The dendritic segments were reconstructed in three dimensions from the serial EMs (Fig. 5h, i). The three-dimensional reconstructed dendritic segments were easily recognized as the same dendritic segments observed in the light microscopy (LM) focus-stacking image because the shape of the dendritic segments with spines was identical to that in the LM image (Fig. 5c, h, i). The spines that elongated in the z-direction could clearly be identified in the 3D-reconstructed image, though they were not easily identifiable by LM^20^. Thus, 3DEM provided accurate images that enabled the quantification of spine counts and the analysis of spine morphology. Spine morphology and dendrite dimensions on the 3D-reconstructed dendritic images were analyzed after identifying the location of the dendritic segments in relation to the LMGs.

**Figure 5.**
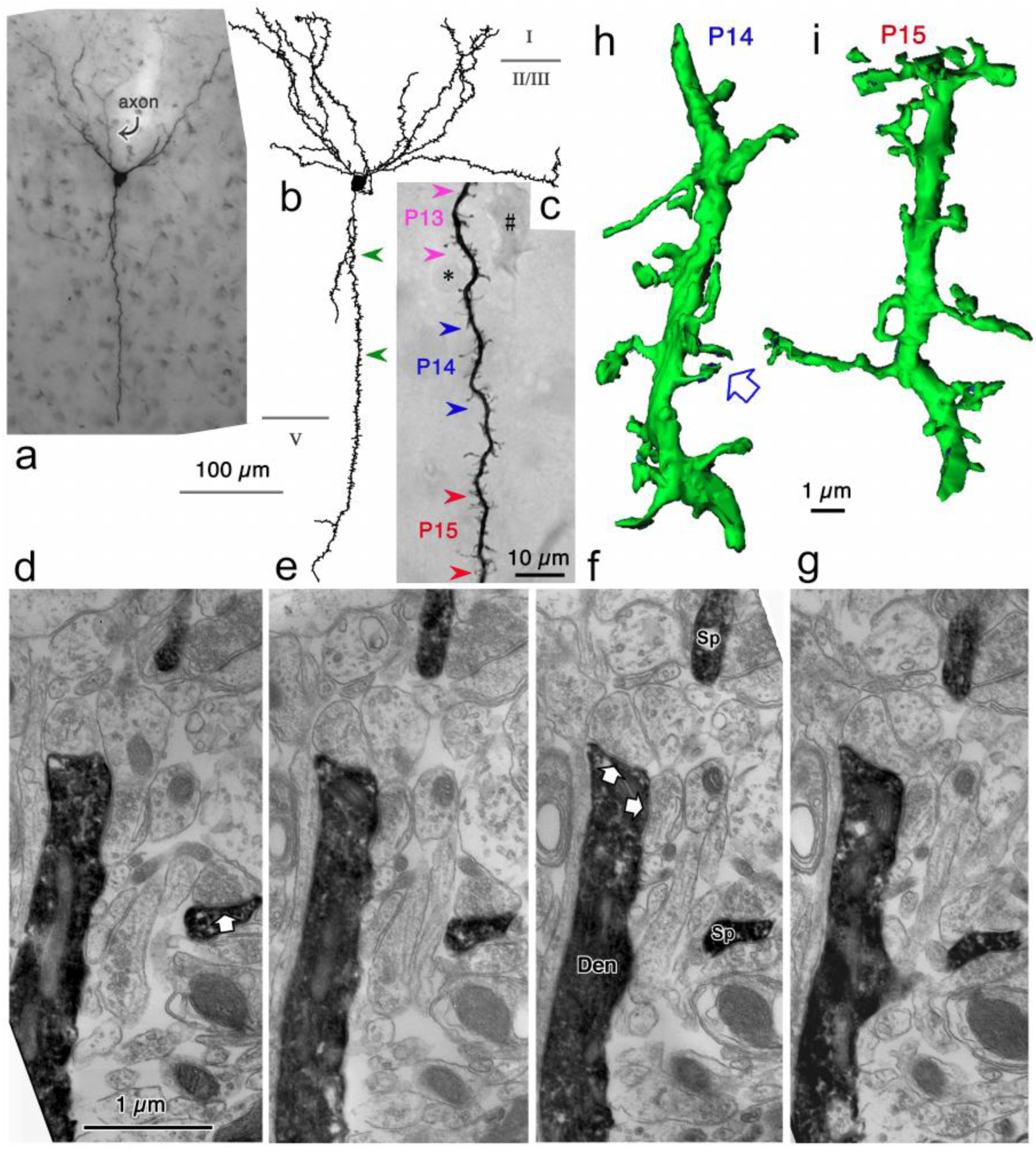
CLEM with Whole Cell Patch Clamp Brain Slice Experiment of a Martinotti Cell in Rat Cortex. **(a)**. DAB-stained Martinotti cell after whole cell patch clamp recording and biocytin injection. **(b)**. Drawing of a Martinotti cell showing the soma and dendrites with spines. **(c)**. An enlarged dendritic segment shown between the green arrowheads in b. The short dendritic segments shown between the two arrowheads were reconstructed in three dimensions from serial thin sections (P13, P14, and P15). The asterisk and hash indicate the soma of glia and degenerated neurons, respectively. **(d–g)**. Serial electron micrographs of the dendritic segment (P14). The spine (Sp) indicated by the white arrow is shown in h. The arrows indicate synaptic contacts. Den: dendritic segment; Sp: spine. **(h)**. Three-dimensional reconstruction of the dendritic segment P14. **(i)**. 3D reconstruction of the dendritic segment P15.

## Discussion

In this study, we developed and demonstrated Meso2EM, a workflow that guides a single neuron recorded by mesoscale functional imaging through confocal microscopy of fixed tissue, micro-CT of a resin-embedded specimen, and block-surface SEM to targeted FIB-SEM. The key methodological feature was not simply combining instruments with different spatial resolutions, but continuously verifying target-cell identity using vascular architecture shared between successive stages. This enabled millimeter-scale functional coordinates in vivo to be transferred to a micrometer-scale target position in the resin block, with the cell body ultimately identified in the FIB-SEM milling face. In two complementary applications, blood vessels, nuclei, cell morphology, and DAB labeling similarly linked an in vivo dendrite to ATUM-SEM and a patch-clamp-recorded cell to serial TEM.

Wide-field two-photon microscopy not only records large numbers of cells simultaneously, but also enables individual neurons to be analyzed within functional networks distributed across multiple cortical areas^1–6^. Such analyses can identify candidate cells for subsequent investigation according to node degree, module membership, or interareal correlation. However, optical information alone cannot determine whether cells with different roles in a functional graph differ in synaptic inputs, intracellular organelles, or local cellular arrangement. Meso2EM supports an experimental design in which a small number of cells are selected according to a functional hypothesis and only their surroundings are acquired at high resolution, rather than converting the entire wide-field image into EM data. It may therefore provide a bridge for future comparisons between structural features and the spatially intermixed functional modules or neuronal populations with different network degrees described by Kiyooka et al.^5^.

In this respect, Meso2EM is complementary to pioneering functional connectomics studies that directly related functional information to connectivity^7,8^. It addresses the targeting problem that grows with imaging field size: which cell should be acquired, and within which EM volume? The present FIB-SEM dataset demonstrates access to the target cell body rather than extensive axonal tracing or complete reconstruction of synaptic connectivity. Accordingly, the current value of this study lies not in completing a connectomic reconstruction, but in establishing the prerequisite targeting route to a functionally identified cell.

The concept of placing micro-CT or X-ray microscopy between light microscopy and EM is supported by substantial prior work. Nondestructive visualization of internal morphology in resin-embedded specimens^14–16^, guidance of regions of interest for volume EM using X-ray images^15^, and integration of in vivo two-photon Ca²⁺ imaging, synchrotron X-ray microtomography, and volume EM in brain tissue^13^ have all been reported. Thus, this study does not claim priority for the use of micro-CT in CLEM.

In the present implementation, the practical role of micro-CT was to retrieve the vascular map contained in the optical images after heavy-metal staining and resin embedding made fluorescence difficult to observe. Vascular lumens were distinguished by their X-ray absorption contrast with the surrounding heavy-metal-containing neural tissue, allowing vascular trajectories in the confocal z-stack to be extended into the block interior. By using laboratory-based micro-CT rather than dedicated synchrotron facilities and exploiting the vasculature itself as a natural three-dimensional fiducial, the search area could be narrowed without introducing additional artificial markers near the target. This accessibility is a practical strength of Meso2EM, although localization performance will depend on vascular density, section orientation, staining uniformity, and micro-CT resolution.

The three applications were not repetitions of the same experiment; they differed in both target and endpoint. The wide-field Ca²⁺ imaging–FIB-SEM application demonstrated cell-body targeting, the in vivo dendrite– ATUM-SEM application demonstrated serial reconstruction of spines and synapses, and the patch-clamp–TEM application demonstrated ultrastructural analysis of selected dendrites from a physiologically identified interneuron. Together, these examples illustrate three transferable design principles: preservation of the imaging plane, acquisition of shared landmarks and progressive narrowing of the search space.

The three applications were not direct comparisons performed under identical conditions, nor were they intended to compare endpoint performance. TEM is suited to high-resolution observation of a limited target; ATUM-SEM provides an archivable serial-section library and a wide XY range; and FIB-SEM enables direct serial acquisition from a selected region within a resin block^24–28^. The practical configuration of Meso2EM should therefore be selected according to the biological question, target size, required isotropy, need for reimaging, and available instrumentation.

The route was completed in approximately two months, including preliminary optimization. Critical decisions were reviewed jointly by several investigators, supporting procedural reproducibility through shared checkpoints and strict adherence to the protocol. Moreover, no target-loss event at the EM stage was encountered in experiments 2 or 3 when their respective pipelines were followed. Because experiments 2 and 3 used different optical inputs and EM endpoints, these observations support the robustness of the shared progressive-targeting principle but do not constitute independent repetitions of the core route. Consultation within one team is also not equivalent to independent-operator validation; additional specimens and independently operating investigators will be required to establish generalizable success rates and quantitative localization errors.

In summary, Meso2EM is a workflow that progressively bridges the spatial and technical gap between single-cell-resolution mesoscale functional images and targeted EM using multiple intermediate landmarks. The present work does not constitute a complete whole-brain functional connectome; rather, it provides a foundation for transferring specific neurons selected in wide-field functional images to restricted EM acquisition volumes. Quantification and automation of this framework, followed by its application to structural comparisons among functionally defined cell populations, should enable mesoscale networks and local ultrastructure to be related in the same cells.

## Methods

### 1) CLEM: Wide-field functional imaging–micro-CT–FIB-SEM (core Meso2EM route)

#### Animals

All animal experiments were performed in accordance with the institutional guidelines and were approved by the Animal Experiment Committee at RIKEN. Both male and female Ai9 mice on a C57BL/6J background (Jackson Laboratory, stock# 007909, Bar Harbor, ME)^29^ were used. In all experiments, mice were housed in a 12-hour light/12-hour dark cycle environment with ad libitum access to food and water.

#### Adeno-associated virus (AAV) vector preparation

G-CaMP7.09^30^ was subcloned into the synapsin I (SynI)-expressing vector from a pN1-G-CaMP7.09 vector construct. pAAV-CIBN-CreC and pAAV-CRY2-CreN were gifts from Michael Bruchas (Addgene plasmid # 75267; http://n2t.net/addgene:75267; RRID: Addgene_75267, Addgene plasmid # 75268 http://n2t.net/addgene:75268; RRID: Addgene_75268, respectively). The following adeno-associated viruses (AAVs) were produced as described previously^31^: AAV-DJ-Syn-G-CaMP7.09-WPRE, AAV-DJ-CIBN-CreC, and AAV-DJ-CRY2-CreN.

#### Postnatal AAV injection and surgery for open-skull cranial windows

AAV injection and craniotomy (surgery) for two-photon imaging were performed using previously described protocols^4,5,32^, with slight modifications to the viral injection part. AAV-DJ-Syn-G-CaMP7.09-WPRE, AAV-DJ-CIBN-CreC, and AAV-DJ-CRY2-CreN were mixed and used after being diluted with 1x PBS (phosphate buffer saline) to the final titer of 4.0 × 10^12^ vg/mL, 4.0 × 10^11^ vg/mL, and 4.0 × 10^11^ vg/mL, respectively. The total volume of the injected AAV solution was 4 µL. During the surgical procedure for open-skull cranial windows, PA-Cre was photoactivated with a white LED.

Cranial window implantation surgery was performed following the previously described method^32^. Briefly, mice (14-20 weeks old) were anesthetized with isoflurane (2%). Once the mice failed to respond to stimuli, we administered hypodermic injections of the combination agent^33^-Medetomidine/Midazolam/Butorphanol (MMB) at a dose of 5 mL/kg body weight for the procedure. The MMB solution consisted of medetomidine (0.12 mg/kg), midazolam (0.32 mg/kg), butorphanol (0.4 mg/kg), and saline. After anesthesia with MMB, the head was shaved and placed in head holders (SG-4N, NARISHIGE). During surgery, their body temperatures were maintained at 36-37°C using a feedback-controlled heat pad (BWT-100, Bio Research Center), and their eyes were coated with an ointment (Neo-Medrol EE Ointment, Pfizer Inc.). After removing the scalp, a 4.5-mm diameter craniotomy was performed over an area that included the primary somatosensory area of the right hemisphere. The craniotomy was then covered with a “double glass” assembly consisting of a No. 2 cover glass (4.5 mm in diameter) and a No. 1 cover glass (6 mm in diameter) (both from Matsunami Glass Ind.) and sealed with dental cement (Super Bond, Sun Medical). The exposed skull was covered with dental cement. After surgery, the mice were treated with a pesticidine-reversing agent, atipamezole hydrochloride (ANTISEDAN, Zoetis Inc.) solution at a dose of 0.12 mg/kg. The mice were then placed on a heating pad to recover.

#### In vivo two-photon calcium imaging of a large field of view

In vivo two-photon imaging was performed using a custom-designed wide-field two-photon laser scanning microscope (FASHIO-2PM)^4^. The mice were fixed in a custom-made stage box (ExPP Co., Ltd.) by firmly screwing the headplate to the stage, and then covered with an enclosure to maintain body temperature. The floor of the stage box was covered with the same bedding as that of the home cage. Before an imaging session, the imaging window was cleaned with a cotton swab soaked in acetone. Fluorescence was observed in the somas of layer 2/3 neurons (120 - 150 µm below the surface). G-CaMP7.09 was excited at 920 nm using a tunable Ti:Sa laser (Maitai DeepSee, Spectra Physics), with laser power set to 60-80 mW at the front of the objective lens. The fluorescence of G-CaMP7.09 was detected with a GaAsP PMT in the range of 515 - 565 nm. All imaging sessions were performed at 3.85 or 7.65 frames/s with 2,048 × 2,048 pixels using custom-built software (Falcon, Nikon) and were saved as 16-bit monochrome TIFF files.

#### Tangential brain slice preparation

Mice were deeply anesthetized by intraperitoneal injection of MMB and transcardially perfused with a pre-fixative solution containing 8.5% sucrose, 5 mM MgCl₂, and 0.02 M phosphate buffer (PB), followed by fixation with 0.5% glutaraldehyde, 0.2% picric acid, and 4% paraformaldehyde (PFA) in 0.1 M PB. Brains were post-fixed in situ for 1 hour at room temperature, then carefully removed from the skull. The tissue was embedded in 4% low-melting-point agarose dissolved in PBS, and surface-tangential brain slices parallel to the in vivo imaging plane were prepared using a vibratome (VT1200S, Leica). Prepared slices were stored at −30°C in cryoprotectant solution until further use.

#### CLEM with microCT and FIB-SEM

The tissue slices were counterstained in fluorescence for 2 hours at 4°C with 4’,6-diamidino-2-phenylindole (5 µg/ml; DAPI; D9542, sigma) and DyLight 649-labeled *Lycopersicon esculentum* (tomato) lectin (20 µg/ml; DL-1178, Vector Laboratories) in 0.05M TBS containing 1% BSA. DAPI and DyLight649 signals visualized nuclei and blood vessels, respectively, and served as landmarks for the subsequent CLEM. The brain sections were mounted upside down on a glass-bottom dish with ProLong Gold Antifade Mountant (P10144, Thermo Fisher Scientific) and observed under an inverted confocal microscope (Nikon Ti2 AX) using a 4× objective lens (Plan Apo λD 4× OFN25, NA 0.20, Nikon) and a 40× silicone-immersion objective lens (CFI Plan Apo Lambda S 40XC Sil, NA 1.25, Nikon). Image stacks were acquired in galvano scanning mode using the 40× silicone-immersion objective lens. Images were collected with a pinhole size of 1.0 Airy unit and a zoom factor of 2. For each imaging field, 2 × 2 tiled images were acquired, with each tile consisting of 1024 × 1024 pixels, and stitched to generate a final image of 1895 × 1895 pixels. The resulting pixel size in the X–Y plane was 0.2158 µm per pixel. Z-stack images were acquired with a step size of 0.5 µm. DAPI, GCaMP, tdTomato, and DyLight 649 signals were sequentially excited using 405-, 488-, 561-, and 640-nm laser lines, respectively. Emitted fluorescence was detected through spectral windows of 429–474 nm (DAPI), 499–525 nm (GCaMP), 571–625 nm (tdTomato), and 662–737 nm (DyLight 649). Images were acquired at 12-bit depth.

After confocal microscopy, the tissue was processed for EM observation as previously reported (Kubota et al., 2018), with slight modifications. The following procedures were performed at room temperature unless otherwise stated. The tissue was first timed using a knife and washed with 0.1 M PB, followed by washes with 0.1 M cacodylate buffer (pH 7.4). The section was then postfixed in 1.5% potassium ferrocyanide and 2% osmium tetroxide (OsO_4_) in 0.1 M cacodylate buffer at 4°C for 1 hour. After post-fixation, the tissue was washed in ultrapure water (Milli-Q Reference water purification system, Merck Millipore, Burlington, MA) and subsequently stained with 1% thiocarbohydrazide for 30 min, followed by washes with ultrapure water. The section was again postfixed in 2% OsO_4_ for 2 hours and, after washing with ultrapure water, stained overnight at 4°C with 1% uranyl acetate. The tissue washed with ultrapure water was processed with modified Walton’s en bloc lead aspartate staining^34^. Lead aspartate solution was prepared by dissolving 0.066 g of lead nitrate in 10 mL of 0.03 M aspartic acid, adjusting the pH to 5.0 with 1 N potassium hydroxide, and heating to 50°C until dissolved. The tissue was stained with the lead aspartate solution at 50°C for 1 hour, followed by washes with ultrapure water. The section was then dehydrated through graded ethanol dilutions and embedded on silicon-coated glass slides in epoxy resin (Hard Plus Resin 812, Electron Microscopy Sciences, Hatfield, PA). The sample was polymerized at 70°C for 3 days.

For CLEM analysis, the resin-embedded tissue was trimmed and imaged using micro-computed tomography (ScanXmate-E090S, Comscantecno Co., Ltd.) to visualize the internal three-dimensional structure of the sample at a voxel resolution of 6.011 µm × 6.011 µm × 6.011 µm (X, Y, and Z). Based on the micro-CT dataset, regions of interest corresponding to the optical imaging planes were identified within the embedded tissue block. The surface of the tissue block was then imaged using a scanning electron microscope (SEM; JSM-IT800, JEOL) to align the internal micro-CT volume with the block surface.

After selecting the target position based on this alignment, focused ion beam–scanning electron microscopy (FIB-SEM, Helios 5 UX, FEI Electron Optics, Eindhoven, The Netherlands) imaging was performed. The sample on the resin block was mounted horizontally on a stub and coated with a 25 nm layer of osmium using an osmium coater (HPC-20, Vacuum Device Inc., Mito, Japan), and graphite paint (Ted Pella, Redding, CA) was applied to enhance electric conductivity. A protective platinum layer with a thickness of 1 mm was deposited on the surface orthogonal to the block, where the ion beam is first incident, to protect the sample from ion beam damage. Trenches were milled on both sides of the region of interest to minimize re-deposition of milled material during automated milling and imaging. Gallium ion beam milling was carried out at an acceleration voltage of 30 kV and a current of 9.9 nA, stage tilt of 52°, and working distance of 4 mm. At each step, 3-10 mm of the block face was removed by the ion beam. Each newly milled block face was imaged with the Through the Lens Detector (TLD) for backscattered electrons at an acceleration voltage of 1.5 kV, beam current of 0.8 nA, stage tilt of 52°, and working distance of 4.1 mm. The pixel resolution was 25.3 nm with a dwell time of 80 ms per pixel. Pixel dimensions of the recorded image were 1536 × 1024 pixels.

### 2) CLEM: in vivo imaging, confocal laser scanning microscopy, and array tomography with scanning electron microscopy

#### Experimental Model and Subject Details

It was a part of experiment for our previous paper^17^. Eight week-old Thy1-eGFP-M mouse on a C57BL/6J background^35^ mice (Jackson Laboratory, Bar Harbor, ME) of male was used. All animal care and use were in accordance with the guideline of the Institutional Animal Care and Use Committee of the National Institutes for Natural Sciences. Mice were housed individually on a 12:12 light-dark cycle. All efforts were made to minimize animal suffering and the number of animals used.

#### Method Details

##### Surgery for open-skull cranial windows

Mouse was anaesthetized with 1–1.5% isoflurane followed by intraperitoneal injection of glycerol (0.6 g/kg) and intramuscular injection of dexamethasone (1 mg/kg). The skull was exposed and a custom-made head-fixing metal chamber was attached with Super-Bond adhesive resin cement (Sun Medical, Shiga, Japan). The center location of cranial windows was determined by using stereotaxic coordinates in accordance with previous studies (AP = +1.3 mm from bregma; ML = 1.2 mm from midline)^36–38^. Craniotomy was bilaterally performed with a stainless-steel trephine drill (Meisinger, Neuss, Germany). The double-layer glass windows were constructed with a large-diameter coverslip (No. 0, 2.3-mm diameter, Matsunami Glass, Osaka, Japan) and a small-diameter coverslip (No. 3, 2.0-mm diameter, Matsunami Glass) by using an ultraviolet curable optical glue (NOA-61, Norland Products, NJ). The double-layer glasses were placed bilaterally on the dura and sealed with resin cement.

##### Observation in vivo under a two-photon microscope

Mice were anaesthetized with inhalation of 0.8–1% isoflurane, and head-fixed on a MAG-2 head-holding device (Narishige, Tokyo, Japan) modified for our custom-made chamber. The bilateral M1s were then imaged under an upright-type Leica TCS SP8 microscope (Leica Microsystems, Wetzlar, Germany) equipped with an InSight DeepSee laser system (Spectra-Physics KK, Tokyo, Japan). GFP signal was excited with a laser of 900-nm wavelength, and detected through a 525–550-nm filter using a 25× water-immersion objective lens (HCX IRAPO L25×, numerical aperture [NA] = 0.95). Dendritic arborization was roughly imaged with a zoom factor at 0.75 (2048 × 2048 pixels, 0.29 × 0.29 µm^2^/pixel; Z-step size, 0.85 µm), and dendritic segments derived from somata less than 400 µm-deep from the pial surface (assumed to be layer 2/3 pyramidal cells) were excluded from the present analysis. High-resolution image stacks of dendritic segments were then acquired at a zoom factor of 10 (512 × 512 pixels, 0.087 × 0.087 µm^2^/pixel; Z-step size, 0.85 µm, typically 20–30 optical sections for each). Subsequent repetitive imagings were acquired with the aid of blood vessels and dendritic-branching patterns. The dendritic segments imaged were mostly located within a depth of 40 µm from the pial surface.

##### Post hoc quadruple immunohistochemistry

Immediately after the two-photon imaging at training day 8, mice were deeply anaesthetized with isoflurane, and transcardially perfused with 5–10 ml of a solution containing 250 mM sucrose, 5 mM MgCl_2_ in 0.02 M phosphate buffer (PB; pH 7.4), followed by 30 ml of 4% paraformaldehyde containing 0.2% picric acid in 0.1 M PB. Brains were then removed and postfixed for 1 hr at room temperature with the same fixative. The brains were cut into 30-µm-thick sections tangentially to the motor area on a vibrating microtome (VT1200S, Leica Microsystems). Tissue sections were stored at −30 °C in cryoprotectant solution (30% glycerol, 30% ethylene glycol in 0.04 M phosphate-buffered 0.9% saline [PBS])^39^ until use.

##### Confocal laser scanning microscopy and image deconvolution

The brain sections were observed under a TCS SP8 confocal microscope (Leica Microsystems) equipped with HyD detectors. For reidentification of the dendritic branches observed in vivo, image stacks were captured using a 25× water-immersion objective lens (HCX IRAPO L25×, numerical aperture [NA] = 0.95) with the pinhole at 1.0 Airy disk unit (diameter of 48.1 µm) and zoom factor at 0.75 (620 × 620 µm, 2048 × 2048 pixels, 0.303 × 0.303 µm^2^/pixel in the X and Y directions; 0.445-µm interval in the Z direction). Alexa Fluor 488 signal was excited with a laser beam of 488-nm wavelength, and observed through 500–550 nm emission prism windows. After the dendritic segments observed in vivo were reidentified, high-magnification image stacks were acquired using 63× oil-immersion lens (HC PL APO CS2 63×, NA = 1.4) with a pinhole at 0.5 Airy disk unit (diameter of 50.1 µm) and a zoom factor at 3.5 (50.21 × 50.21 µm, 1024 × 1024 pixels, 0.049 × 0.049 µm^2^/pixel in the X and Y directions; 0.129-µm interval in the Z direction). CF405S, Alexa Fluor 488, 568, or 647 was excited with a 405, 488, 552, or 638 nm laser beam and sequentially observed through an emission prism window of 425–475, 500–550, 580–630, or 650–720 nm, respectively, in the photon-counting mode.

The acquired image stacks were deconvolved with HyVolution 2 (Leica Microsystems) with the following parameters: microscopic type, confocal; back projected pinhole diameter, 143 nm; lens objective NA, 1.4; lens immersion refractive index, 1.518; medium refractive index, 1.515; excitation wavelength, 405, 488, 552, or 638 nm; emission wavelength, 431, 517, 603, or 665 nm; sampling X, Y and Z intervals, 49.082, 49.082 and 129.2 nm; solid brick mode; using theoretical point spread function; maximum iteration number, 30; signal-to-noise ratio, 15; quality change threshold, 1 × 10^−4^.

All the presented figures were edited on a graphics software, Canvas Draw (Canvas GFX, Inc., Plantation, FL). Image appearance in figures was optimized uniformly in individual panels for presentation such as brightness/contrast adjustment and Gaussian filtering. The low-magnification confocal images of dendritic segments are shown in two-dimensional projections of 3D image stacks, while the images that show spines, presynaptic and postsynaptic puncta at high magnification are presented in single focused planes.

##### Somatodendritic reconstruction of GFP-labeled cells in Thy1-eGFP-M mice

Thy1-eGFP-M mice were transcardially perfused with 5–10 ml of a solution containing 250 mM sucrose, 5 mM MgCl_2_ in 0.02 M PB (pH 7.4), followed by 30 ml of 4% paraformaldehyde containing 0.2% picric acid and 1% glutaraldehyde in 0.1 M PB. After postfixation in body for 1 hr at room temperature, the brain tissues were sectioned into ∼150-µm-thick slices tangentially to the cortical surface of M1 on a vibrating microtome. The tissue was stored at −30 °C in cryoprotectant solution until use.

The section was counterstained in fluorescence for 2 hrs at 4 °C with 5 µg/ml of 4’,6-diamidino-2-phenylindole (DAPI; 10236276001, Roche, Basel, Switzerland) and 20 µg/ml of DyLight 594-labeld Lycopersicon Esculentum (Tomato) Lectin (DL-1177, Vector Laboratories, Burlingame, CA) in 0.05M TBS containing 1% BSA. DAPI and DyLight 594 signals visualized nuclei and blood vessels, respectively, in different fluorescence from GFP for landmarks of the following CLEM. The section was mounted upside-down on a glass-bottom dish with SlowFade™ Gold Antifade Mountant and observed under an inverted-type TCS SP8 confocal microscope. Image stacks were obtained using a 25× water-immersion objective lens (HCX IRAPO L25×, NA = 0.95) with the pinhole at 1.0 Airy disk unit (diameter of 48.1 µm) and zoom factor at 0.75 (620 × 620 µm, 2048 × 2048 pixels, 0.303 × 0.303 µm^2^/pixel in the X and Y directions; 0.502-µm interval in the Z direction). DAPI, GFP and DyLight 594 signals were excited with 405, 488 and 552 nm laser beams and sequentially observed through emission prism windows of 410–489, 500–550 and 600–700 nm, respectively.

After the confocal microscopy, the tissue was processed for EM observation as previously reported ^54^ with slight modification. The following procedures were performed at room temperature unless otherwise stated. The tissue was washed with 0.1 M PB followed by washes with 0.1 M cacodylate buffer (pH 7.4). The section was then postfixed in 1.5% potassium ferrocyanide, 2% osmium tetroxide (OsO_4_) in 0.1M cacodylate buffer at 4 °C for 1 hr. After the postfixation, the tissue was washed in ultrapure water (Milli-Q® Reference water purification system, Merck Millipore, Burlington, MA), and subsequently stained with 1% thiocarbohydrazide for 20 min, followed by washes with ultrapure water. The section was again postfixed in 2% OsO_4_ for 30 min and, after washed with ultrapure water, stained overnight at 4 °C with 1% uranyl acetate. The tissue washed with ultrapure water was processed with modified Walton’s en bloc lead aspartate staining^34,40,41^. Lead aspartate solution was prepared by dissolving 0.066 g of lead nitrate in 10 ml of 0.03 M aspartic acid, pH adjusted to 5.0 with 1N potassium hydroxide and kept at 50 °C until dissolved. The tissue was stained with the lead aspartate solution at 50 °C for 2 hrs, followed by washes with ultrapure water. The section was then dehydrated in graded dilutions of ethanol and embedded on silicon-coated glass slides in epoxy resin (Durcupan ACM; Sigma-Aldrich, St. Louis, MO). The sample was polymerized at 70 °C for 3 days.

The embedded tissue was serially re-sectioned into 50-nm-thick ultrathin sections and collected using an automated tape collecting ultramicrotome (ATUMtome; Boeckeler Instruments Inc., Tucson, AZ) on a plasma-hydrophilized carbon nanotube-coated polyethylene terephthalate tape (CNT-PET tape)^24^. The following SEM observation was described previously^24^. Briefly, serial ultrathin sections on tape were cut into strips and mounted in order on 4-inch silicon wafers with double-sided adhesive conductive tape. Conductive surface of the CNT-PET tape was grounded to the wafer with copper foil tape. The serial EM images were obtained using a backscattered-electron detector (BSD) of field emission (FE) SEM (Sigma, Carl-Zeiss Microscopy, Oberkochen, Germany) with a guide of Atlas 5 (Fibics incorporated, Ottawa, Canada). EM images were captured at a resolution of 5 × 5 nm^2^/pixel in the X and Y directions.

We obtained EM image stacks of two adjacent regions separately from one brain sample (approximately 200 × 200 µm in XY, 662 sections of 50-nm thickness [totally 33.1 µm in depth]; 100 × 150 µm in XY, 496 sections [totally 24.8 µm in depth]). Tiled images in a single plane were stitched, and serial mosaic images were aligned on TrakEM2 (https://imagej.net/TrakEM2). The 3D EM images were downscaled to a resolution of 10 × 10 nm^2^/pixel in the XY direction, and imported to the segmentation software VAST Lite (https://software.rc.fas.harvard.edu/lichtman/vast/). After the dendritic segments observed in the two-photon images were reidentified in the confocal fluorescence images, nuclei and blood vessels labeled in fluorescence were found in the EM image stacks. Based on such landmarks, the dendritic segments were distinguished from surrounding neuronal and non-neuronal microstructures, and manually reconstructed on the VAST Lite software. Spine and bouton volumes of reconstructed data from the EM images were measured on the VastTools MatLab scripts for VAST Lite.

### 3) CLEM: whole cell patch clamp brain slice experiment and transmission electron microscopy

#### 1. Light microscopy

We injected biocytin into non-pyramidal cells using whole-cell recordings with a 300-µm-thick slice of rat frontal cortex. Previous papers have described this method in detail^19,42^. Subsequently they were processed histologically for EM. Protocol for the histological treatment was described previously^43^. Briefly, the 300 µm thickness slices were fixed with 4% paraformaldehyde, 0.2% picric acid, 0.1% glutaraldehyde in 0.1M PB with briefly in microwave^44^. They were re-sectioned into 50 µm thick sections after embedded in agar. The recorded cells were stained with DAB and dehydrated. They were embedded flat in Epon between silicon (Sigmacoat, Sigma-Aldrich, St. Louis, U.S.A.) coated glass slide and cover slip. The tissue block shrank evenly in about 90% during the resin-embedding process^18^. Dendrites/axons of the stained cells were reconstructed using Neurolucida (MicroBrightField, Williston, VT, U.S.A.) to obtain their dendrogram and drawing (Figure 5). The dendritic segments for 3DSEM were selected at ∼50 µm intervals along dendritic branches lacking severed endings on the dendrograms. Some dendritic segments included bifurcation points were also selected (Fig 5, Extended Fig. 4). Mainly two dendritic trees were selected for the analysis. The stained cells were photographed with light microscope at 0.5 µm focus steps using a 100x objective. (Fig 5, Extended Fig. 4). Almost entire parts of the two dendritic trees were photographed in 6 image portions. The serial images at 0.5 µm focus step were stored to layers of a Photoshop file successively and dendritic segments for 3DSEM were marked on new layers between the serial LM images as a guide for EM.

Focus stacking image was obtained using ‘auto-blend layers/stack images’ function of Photoshop (Adobe, San Jose, U.S.A.), which combine the best focused area of the multiple focus step LMgs, to give a greater depth of field of the dendritic segment. This image is used for evaluation of the 3D reconstructed segment from the serial EMgs as the same dendritic segment selected by LM.

#### Ultrathin section

The silicone coated cover slip was removed and a block contained the stained cell was cut out with a blade. Then the block was glued on Epon column with super bond (Krazy Glue, Westerville, U.S.A. or Aron Alpha, Toagosei, Tokyo, Japan). The block was trimmed in a trapezoid and most dendritic branches of the cell were left in the trimmed block. The cell was then serially sectioned into 50 nm thick ultrathin sections using an ultramicrotome (Reichert Ultracut S, Leica Microsystems, Wetzlar, Germany). The ultrathin serial sections were mounted on formvar-coated single-slot grids (NOTCH-NUM Grids, 1 x 2mm slot, SynapTek) and were stained with lead citrate.

#### 3. EM observation

First emerged dendritic segment in the serial ultrathin sections could be predicted by careful observation of the Photoshop file with the serial LMgs. Which ultrathin section and where in the section does the dendritic segment found? We estimated an approximate existence of the segment in the ultrathin sections under EM observation with careful reference of the serial LMgs. We carefully compared the dendritic segments shown in LMgs and an candidate dendritic segments found in EMgs during observation using the SEM with refering the landmarks such as capillaries and electron-dense DAB-stained dendrites and spines (Extended Figures 4). Once the dendritic segment was found, serial EMgs with the segment and the associated structure were captured using a CCD camera (XR-41, Advanced Microscopy Techniques, U.S.A.) equipped in a Hitachi H-7000/H7700 TEM without tilting angle. Additional EMgs were captured using tilting of up to 60° to get a clear image of a synapse contact in right angle for cleft structure^21^. Thickness of the ultra-thin sections was calibrated by a color laser 3D profile microscope (VK-9500; Keyence, Japan)^21^.

#### 4. 3D reconstruction

The serial EMgs were aligned manually using free 3D reconstruction software, Reconstruct (http://synapses.clm.utexas.edu/tools/index.stm)^45^. Segmentation of the DAB-stained dendritic segments and the associated neuronal structures were done manually using the Reconstruct. The reconstructed 3D images were compared with the focus stack images made with serial focus light micrographs to confirm the same dendritic segments (Fig. 5).

## Data and code availability

The imaging data and code supporting the findings of this study are available from the corresponding author upon reasonable request.

## Acknowledgments

The authors would like to thank H. Kita, S. Hatada, and N. Takahashi for technical assistance, A. Gulledge for valuable comments, and R-COMS_CBS for technical support, including electron microscopy sample preparation and imaging by the Support Unit for Electron Microscopy Techniques, Research Resources Division, RIKEN Center for Brain Science.

## Author contributions

Conceptualization: Mitsuo Suga, Yasuo Kawaguchi, Masanori Murayama, Yoshiyuki Kubota

Methodology: Masanori Murayama, Yoshiyuki Kubota

Investigation: Ikumi Oomoto, Motohide Murate, Jaerin Sohn, Maya Odagawa, Masaru Tamura, Sayuri Hatada, Naomi Egawa, Mitsuo Suga, Yasuo Kawaguchi, Yoshiyuki Kubota

Data Curation: Ikumi Oomoto, Motohide Murate, Masaru Tamura, Jaerin Sohn, Sayuri Hatada, Naomi Egawa, Yoshiyuki Kubota

Formal Analysis: Ikumi Oomoto, Motohide Murate, Masaru Tamura, Jaerin Sohn, Sayuri Hatada, Naomi Egawa, Yoshiyuki Kubota

Visualization: Ikumi Oomoto, Yoshiyuki Kubota

Writing – Original Draft: Ikumi Oomoto, Motohide Murate, Masanori Murayama, Yoshiyuki Kubota

Writing – Review & Editing: Ikumi Oomoto, Masanori Murayama, Yoshiyuki Kubota

Supervision: Masanori Murayama, Yoshiyuki Kubota

Project Administration: Masanori Murayama, Yoshiyuki Kubota

Funding Acquisition: Ikumi Oomoto, Masanori Murayama, Yoshiyuki Kubota

## Extended Figures

**Extended Figure 1.**
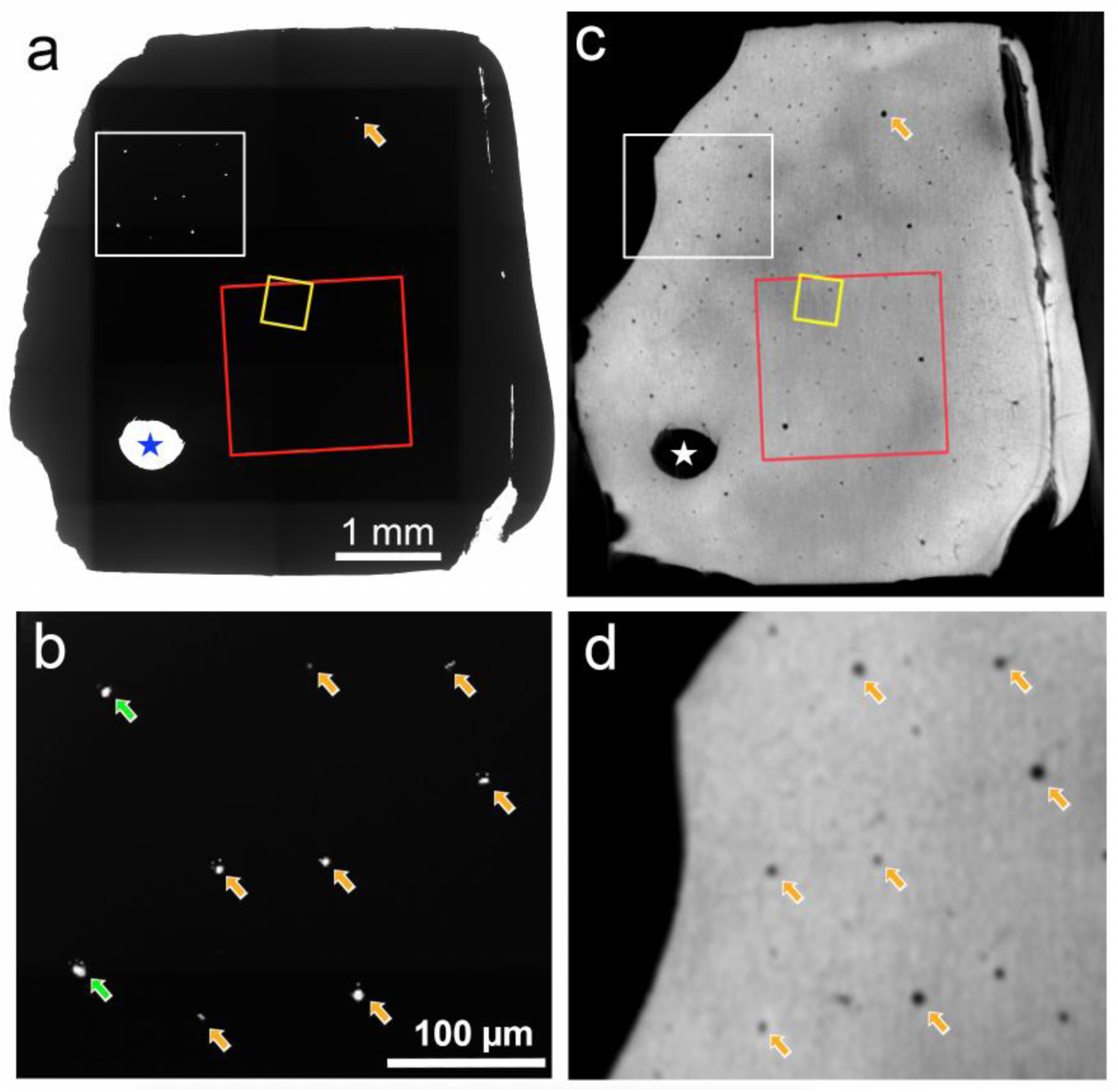
Blood vessels are good landmarks to identify the same portion in micro-CT image and bright-field image of the tangential brain slice after the EM sample preparation with the rOTO protocol. **(a),** Bright-field image of the tangential brain slice shown in Fig. 1m. (**b),** Enlarged micrograph of the white rectangle area in a. (**c),** Micro-CT image shown in Fig. 1j. (**d),** Enlarged image of the white rectangle area in c. Only vertically oriented capillaries (arrows) are visible for the light transmitted under the light microscopy and identical capillaries were found in the micro-CT image (arrows).

**Extended Figure 2.**
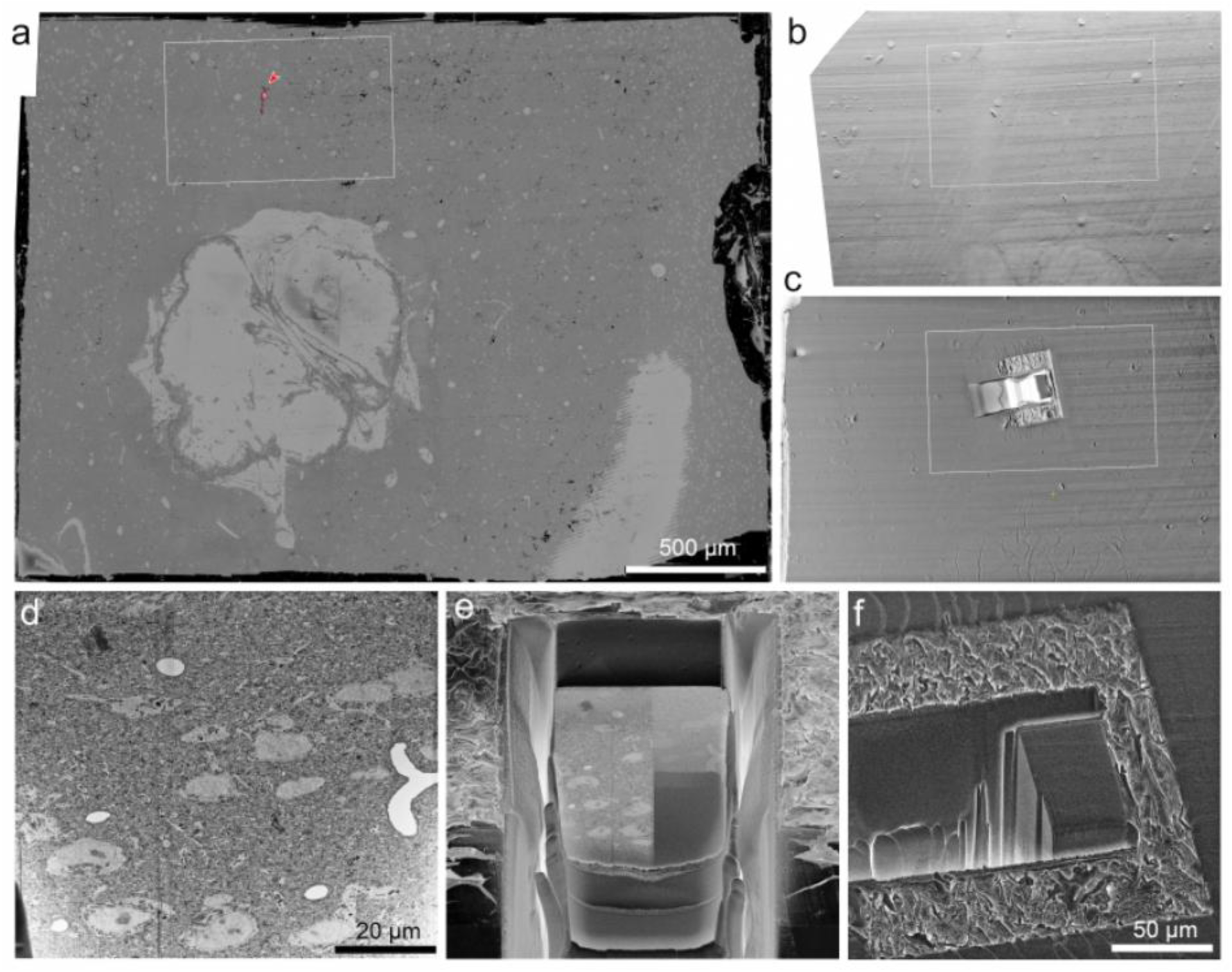
Overview for the FIB-SEM milling. **(a),** A surface image of the brain tissue block captured using SEM before starting FIB-digging. Rectangle area is identical to Fig. 4f. The ROI neuron location was carefully estimated as in the main body. **(b – c),** Surface view of the block under the FIB-SEM before (b) and after (c) starting the FIB-gigging. **(d),** Electron micrograph of the milled surface. **(e, f),** Three-dimensional view of the FIB-SEM digging.

**Extended Figure 3.**
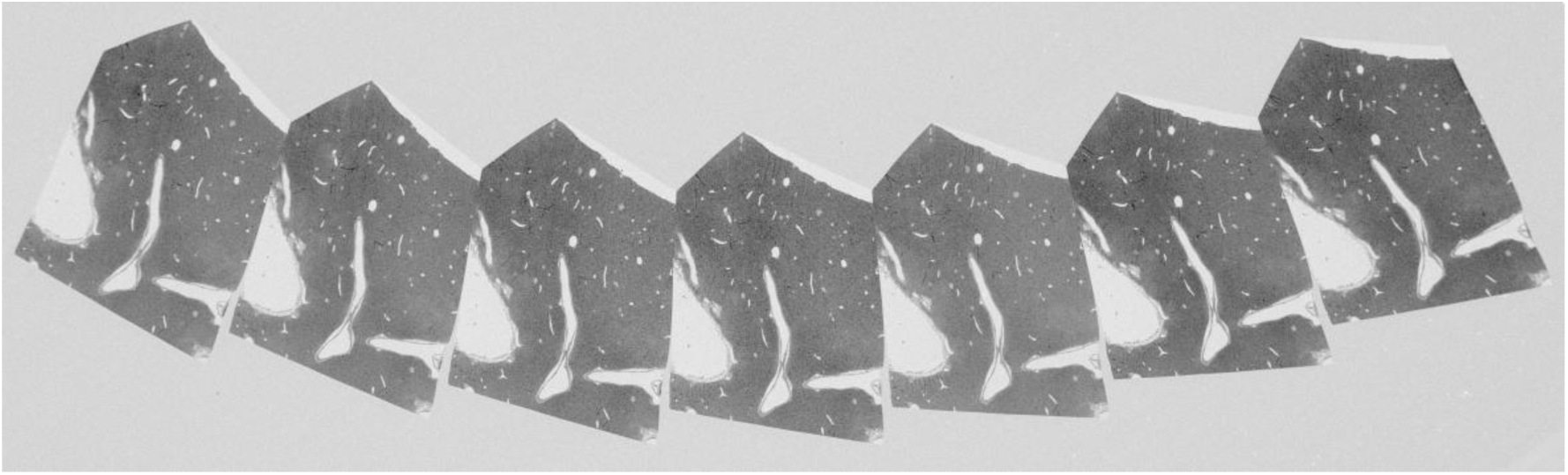
Serial ultrathin sections involving the *in vivo* imaged region shown in Fig. 4l.

**Extended Figure 4.**
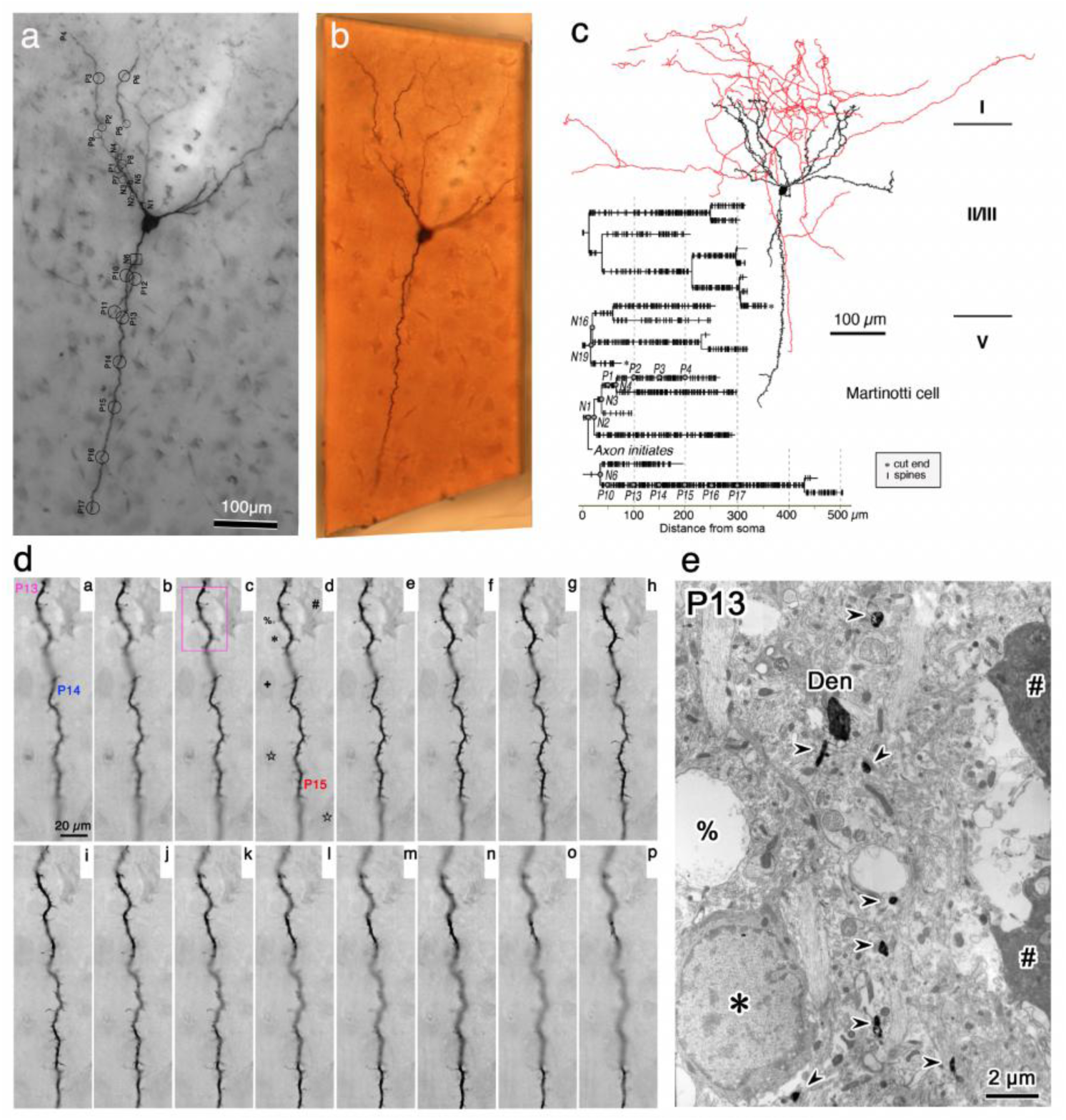
Morphology of a Martinotti cell **(a),** DAB-stained Martinotti cell after whole cell patch clamp recording and biocytin injection. Dendritic segments are marked with the segment name. **(b),** Trimmed block with the Martinotti cell for ultrathin sectioning. **(c),** Axonal and dendritic feature of the Martinotti cell. Axons are shown in red, dendrites and somata are shown in black. Dendrogram showing spine locations with vertical bars and the dendritic segments for the 3D reconstructions from serial electron micrographs are marked with the segment name on the dendrogram. **(d),** Serial LMgs at 0.5-µm z-steps using a 100x objective. (a-p) of the dendritic segment shown in Fig. 1c, including P13, P14 and P15 segments. P13 (pink) and P14 (blue) segments are labeled in LMgs between a and k, and the P15 segment (red) is labeled in LMgs between d and p. Soma (#, *, +), capillary (☆) and vacuole (%) are labeled as landmarks in **d**. Conventional light micrographs at 0.5 µm focus steps using a 100x objective. (a-p) of the dendritic segment shown in Fig. 1c. **(d),** Electron micrograph showing DAB-stained dendrite (Den) and spine flakes (arrowheads). Deteriorated neuron (#), glial soma (*) and vacuole (%) are identical to the structure found in the light micrograph (Fig. 5c, Extended Fig. **d-d**).

